# Teacher Perceptions of Using Robots to Teach Neuroscience in Secondary School

**DOI:** 10.1101/2021.04.01.438071

**Authors:** Claudio C. S. de Freitas, Camden Hanzlick-Burton, Miroslav Nestorovic, Jennifer DeBoer, Gregory J. Gage, Christopher A. Harris

**Affiliations:** School of Engineering Education, Purdue University, USA; Backyard Brains, Inc., Ann Arbor, MI, United States

**Keywords:** neuroscience, neurorobots, neurorobotics, computational neuroscience, education technology, robotics, active learning, secondary school, high school, teachers

## Abstract

The use of robots to help teachers to engage students in STEM subjects has been increasing in recent years, and much progress has been made in K12 schools to incorporate robots in the pedagogy. Recent studies indicate that robots can play a significant role to engage students in neuroscience subjects, but little is known about teachers’ perceptions of using robots to teach neuroscience. In this paper, we present a study based on a survey questionnaire conducted with 84 teachers across multiple high schools in the United States to understand their perceptions about the usefulness of using robots to teach neuroscience. To situate teachers with an example of how robots can be used in neuroscience classrooms, we describe an educational tool called the SpikerBot. Our preliminary results indicate that there is an opportunity for neuroscience-oriented robots in secondary education, provided sufficient on-boarding and training videos.

## Introduction

Understanding the brain is a profound and fascinating challenge, captivating the scientific community and the public alike. The lack of effective treatment for most brain disorders makes training the next generation of neuroscientists, engineers and physicians a key concern. While student laboratories involving biological preps are starting to gain traction within K12 [1], what is missing are technologies and curricula that enable students to develop and use computational models of functioning brains. Neural modeling is becoming an increasingly important area of neuroscience research, and developing the ability to work effectively with neural networks for artificial intelligence applications is increasingly important to students [2], [3].

*Educational neurorobotics* seeks to introduce K12 students to neural modeling with physical neurorobots - robots controlled by computational models of biological neural networks [4]-[7]. Robotics has been shown to be a highly motivating and effective framework for teaching STEM in schools [8]-[10], including to underrepresented students [11]-[13]. Robot-based activities can scaffold multiple disciplines with a positive impact on achievement scores, science concepts, and attitudes [14]-[19]. A review of 315 research articles that used LEGO robotics in educational settings concluded that teamwork and problem-solving are the key educational contributions of LEGO robotics in K12 [20]. National and international robotics competitions for K12 students (e.g. FIRST Robotics), and academic conferences on educational robotics (e.g. Conference on Robotics in Education, IEEE Frontiers in Education Conference) further highlight the widespread use of robots in the classroom. By enabling students to design computational brain models that make robots perform life-like behaviors, educational neurorobotics aims to introduce robot-based labs to the life sciences, allowing students to demonstrate understanding of structure and function within the nervous system (HS-LS1-2) and engineer solutions to real world problems associated with specific neurological conditions (HS-ETS1). However, while neurorobots are increasingly used to conduct neuroscience research in academic settings [21]-[25], little is known about the challenges and opportunities associated with using robots to teach neuroscience in the classroom.

Following the analysis of the existing robots and tools available in the market and opportunities to use technology to teach neuroscience in the classroom, we have developed an open-source neurorobot to help students learn neuroscience through robotics [7]. The “SpikerBot” has a camera, a microphone, a speaker, wheels, Wi-Fi and simulated brains that students control with an app (Figure 1). The app allows students to design neural networks and observe their effect on the robot’s behavior in real-time. To assess the ability of this tool to help teachers demonstrate neuroscience principles in class, we coordinated a 1-week neuroscience module based around our neurorobots in U.S. high schools. We found that the neuroscience module improved high school students’ conceptual understanding of neuroscience and their self-conception as neuroscientists [7]. In light of these preliminary results, we have found an opportunity to expand the application of neurorobots across K12 classrooms. However, for educational neurorobotics to succeed in K12, it is necessary to establish a dialogue across multiple stakeholders to understand their perceptions and needs. Are teachers interested in using robots to teach neuroscience? Can teachers add neuroscience activities, given the limited space for neuroscience in the Next Generation Science Standards (NGSS)? What supports do teachers feel they would need to teach neuroscience with robots? What technology availability and budget considerations need to be addressed? To answer these questions, we conducted a survey of high school teachers across multiple states in the United States, to understand teachers’ perceptions of using robots to teach neuroscience in school.

**Figure 1.**
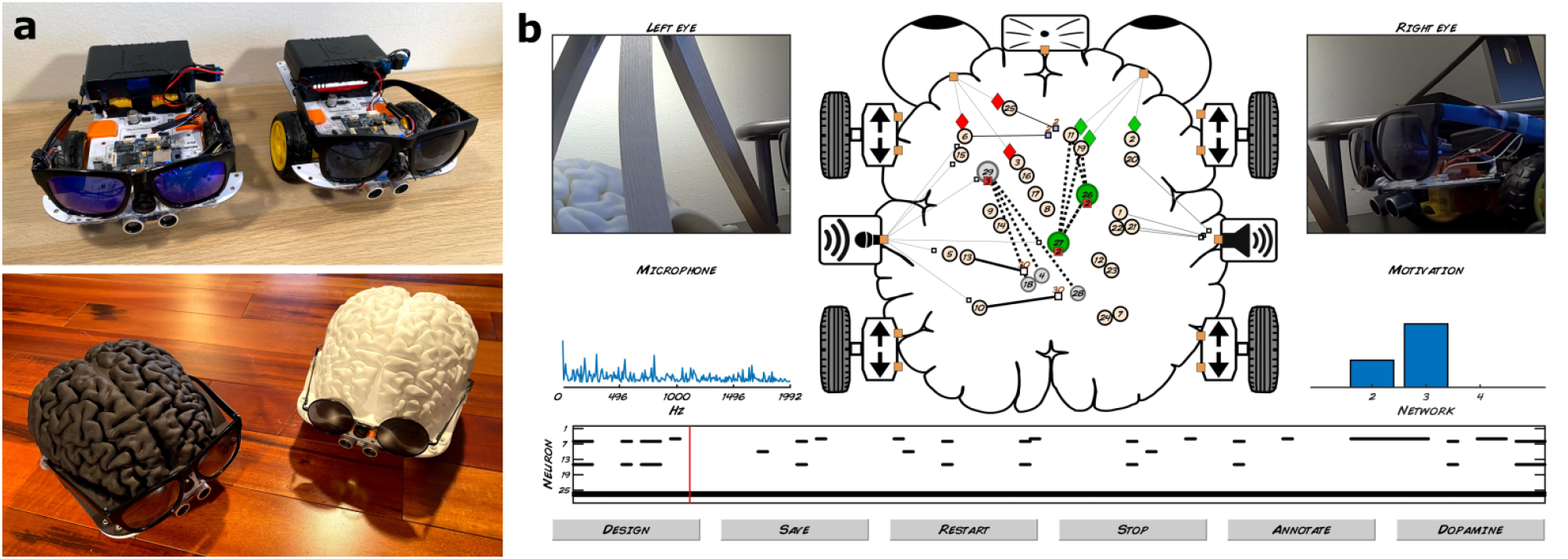
Neurorobots for education. a) Wireless neurorobot prototypes. b) User interface for students shows audio, video and brain activity in real-time.

## Methods

We designed a survey to explore US high school teachers’ perceptions of the idea of using robots to teach neuroscience (Appendix 1). The goal of the survey was to address the following questions:

- Are teachers interested in using robots to teach neuroscience?
- Can teachers add neuroscience activities, given the limited space for neuroscience in the Next Generation Science Standards (NGSS)?
- What supports do teachers feel they would need to teach neuroscience with robots?
- What technology availability and budget considerations need to be addressed?

To facilitate quantitative analysis, all questions were multiple-choice. We also collected demographic information for statistical purposes. The survey development process is described in Figure 2.

**Figure 2.**
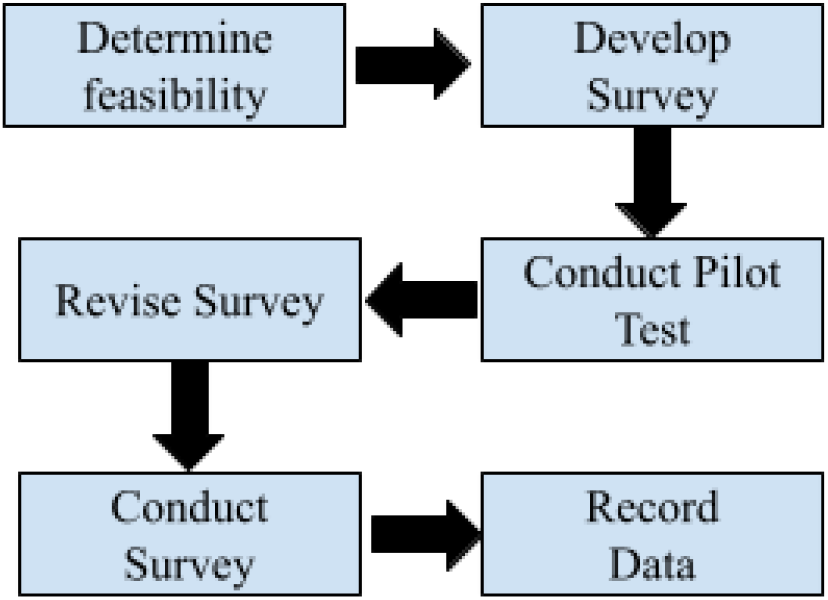
Survey design process

The survey was disseminated using Qualtrics to a pool of K12 teachers within our network. Most of these teachers had provided their email at one of Backyard Brains’ science or education conference booths. All participants were included in a lottery to distribute 10 neuroscience research kits (value: $200 each). We contacted K12 teachers via email from February until March 2021. The email contained a link to a Qualtrics survey about the use of neuroscience and neurorobotics in the classroom.

## Results

The complete list of survey questions and answers can be found in Appendix 1. We received 84 responses to our survey. A majority of participants taught at the high school level. Participants taught a variety of STEM subjects including biology (21% of answers), anatomy and physiology (12%), general science (9%), computer science (7%), and engineering (6%). Most stated that their courses align to the NGSS (45%) or State Science Standards (21%). Participants stated that they used hands-on activities a couple of times per week (51%) or a couple of times per month (23%), and had Chromebooks (26%) or PC/Mac computers (17%) available for students taking their courses.

On the important question “Given the limited space reserved for neuroscience in state-standards, if you wanted to add neuroscience activities to your classroom, could you do it?” 81% of teachers stated they could add neuroscience activities. Of these, 60% felt that neuroscience was well supported by their state standards, while 37% said they could justify it even if not in state standards.

We then asked respondents to watch a 2-minute video about the SpikerBot [26], showing how students use the robot in class. After watching the video, 80% of teachers stated that they would be interested in using the robot to teach neuroscience. Moreover, 81% said that learning to use the SpikerBot (ex. by participating in a workshop) would count towards required professional development hours.

On average, teachers felt that a new robot-based neuroscience curriculum should consist of 5-6 lessons, with 93% of teachers preferring 10 lessons or fewer. When asked which professional development resources would best prepare them for teaching with neurorobots, most wanted teacher training videos (23%) or live virtual training workshops (20%). Finally, 78% of teachers stated that they would need to apply for private donations or a grant to afford a $2500 set of 10 robots for their classroom.

## Discussion

The results of the survey reveal that teachers recognize the value of neuroscience in the classroom and the important role of robots to teach neuroscience. Analysis of responses showed that teachers with different years of experience and courses taught were interested in tools like the SpikerBot to teach neuroscience. Teachers feel they can add neuroscience activities to their teaching, even though neuroscience is a new discipline that has yet to be fully integrated in state standards. Moreover, teachers like the idea of teaching neuroscience with robots, and feel that learning to use a robot such as the SpikerBot would count towards minimum teaching standards related to professional development.

The survey also identifies two challenges. First, it shows that at the price point of typical “smart robots” such as LEGO Mindstorms, only a small number of teachers (13%) would be able to use department/school funds. To ensure broad adoption of neuroscience-oriented robots, it is therefore necessary to reduce the hardware cost of the robot, while also providing teachers with support to write grants. Our interpretation of these results is that government, policymakers, high school boards, and funding agencies need to support schools and teachers to access funding to equip their classroom with technologies that are continuously emerging. To do so, further research needs to build upon findings that demonstrate the benefits of technology to help teachers to engage and motivate students in STEM courses.

Second, capacity building also emerged as a point of attention in our analysis. While most teachers presented significant interest in using technology to teach neuroscience, they also raised concerns with appropriate training to adopt those technologies. Thus, enabling teachers to teach neuroscience with robots will require production of a variety of onboarding, training and support materials. This finding aligns with existing research related to teaching capacity building which calls on school leadership to foster teaching/learning reformation to support educational demands, such as providing effective infrastructure and professional training to help teachers adopt and adapt innovative pedagogies.

Although the survey shows promising results, this study has a couple of limitations. First, the survey participants are familiar with Backyard Brains, and may have used other neuroscience classroom products. They are therefore subject to biases. Second, the internal consistency of the survey was not assessed and the sample size is relatively small. Third, 18% of participants did not complete the entire survey.

Although the survey was used to understand teachers’ perceptions using the SpikerBot, there is a huge opportunity to take our findings and translate them to other neurorobots in the market. Thus, it is recommended that to fully satisfy minimum requirements as a neurorobot in secondary education, robots designed to teach neuroscience should 1) be cost-effective, 2), be compatible with multiple hardware and software, and 3) support teachers with appropriate infrastructure and capacity building programs to reduce the learning curve to adopt and use the tool. In the future, we expect to increase our sample size and increase the research rigor throughout the survey development process.

In terms of opportunities to use robots to teach neuroscience in secondary schools, the positive impact of active learning and robotics is well known in STEM education [27], [28], but if educators who teach neuroscience-related topics are too disconnected from the underlying principles of active learning and robotics in the classroom, this misalignment can lead to important misconceptions regarding the benefits of technology to teach neuroscience using robots. Thus, we argue that the successful delivery of robots to teach neuroscience needs to be grounded in solutions that are affordable to teachers and schools’ budgets, include sufficient capacity building programs that provide enough time and resources to adopt and implement technologies, and implement flexible policies that support innovative pedagogies in the classroom.

## Acknowledgments

GJG is a co-founder and co-owner of Backyard Brains, Inc., a company that manufactures and sells neurorobots. CHB, MN, and CAH are employed by Backyard Brains, Inc. The remaining authors declare that the research was conducted in the absence of any commercial or financial relationships that could be construed as a potential conflict of interest.

Research reported in this publication was supported by a Small Business Innovation Research grant #1R43NS108850 from the National Institute of Neurological Disorders and Stroke of the National Institutes of Health. CAH and GJG are Principal Investigators on this grant.

## Appendix 1

This appendix lists all 17 questions and possible answers in our survey. Answers are presented as bars representing the number of answers we obtained, as well as percentages relative to the total number of answers for each question. For questions with long answers, the text of the answers is given first, followed by results with shortened answers.

## Q1. What is the name of the school(s) where you teach? Please, specify

(data not shown)

## Q2. What is your school’s zip code(s)? Please, specify

(data not shown)

## Q3. What grades do you teach? (select all that apply)

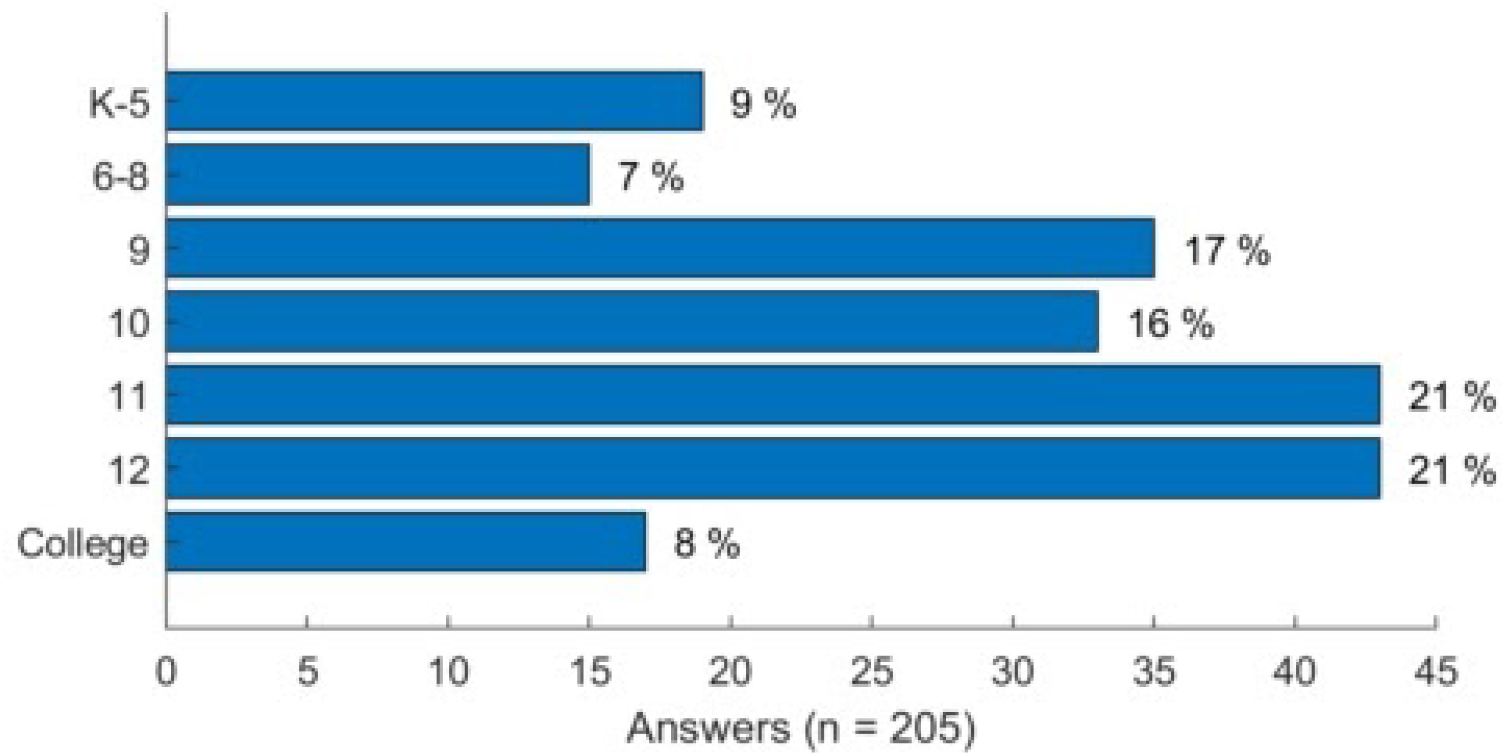

## Q4. What courses do you teach? (select all that apply)

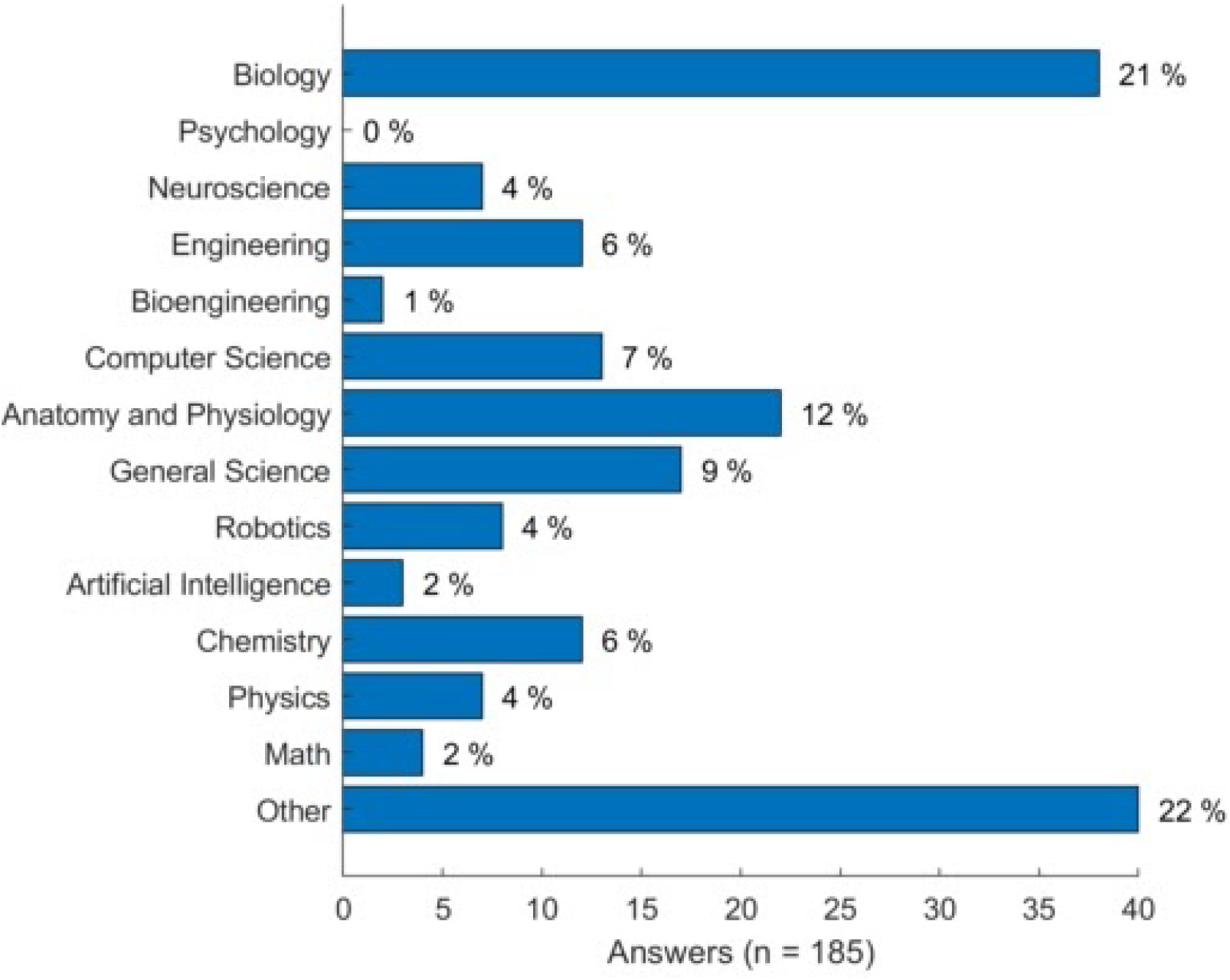

## Q5. What standards do your courses align to? (select all that apply)

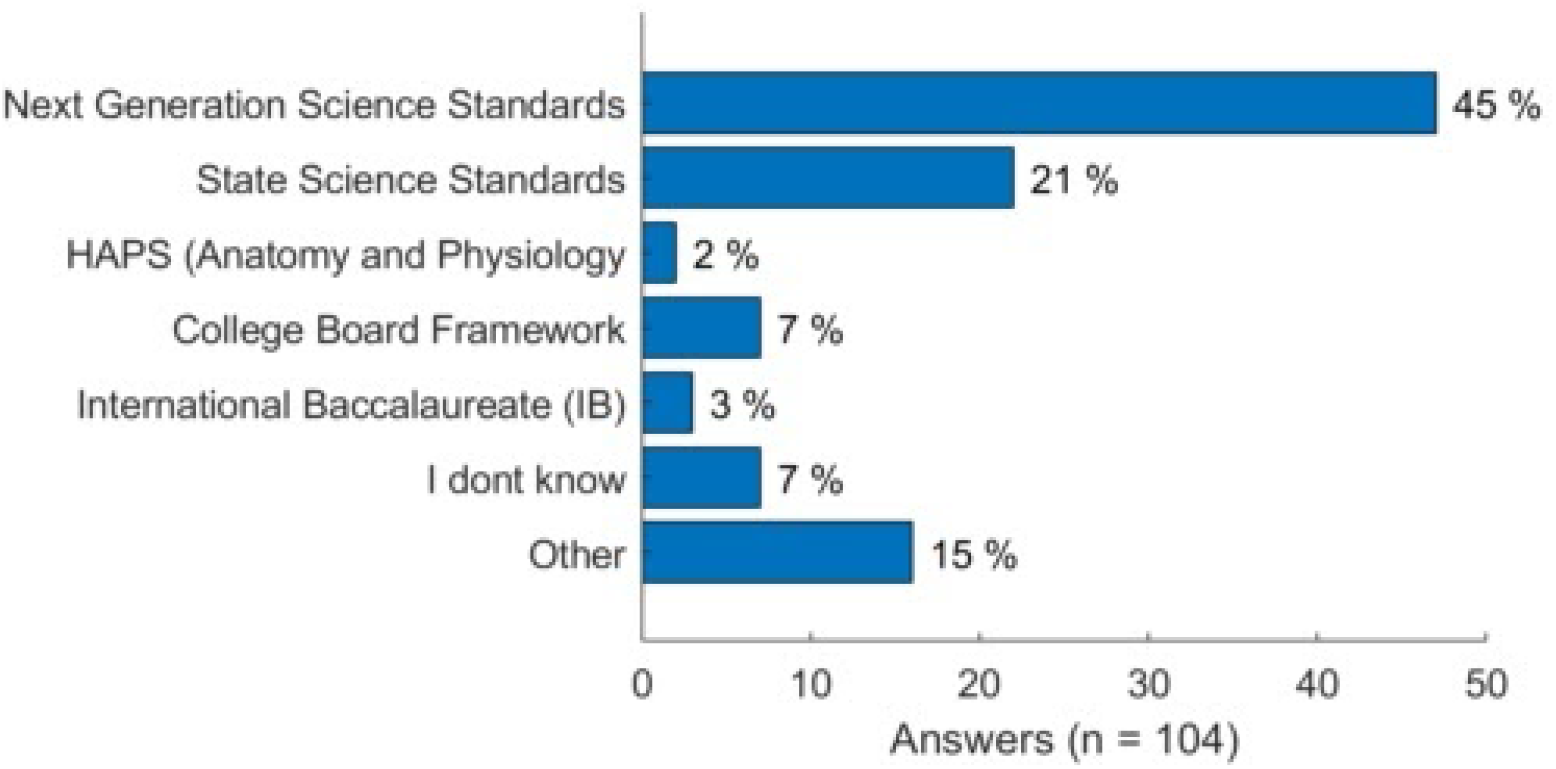

## Q6. Do you currently use hands-on activities in your classes?

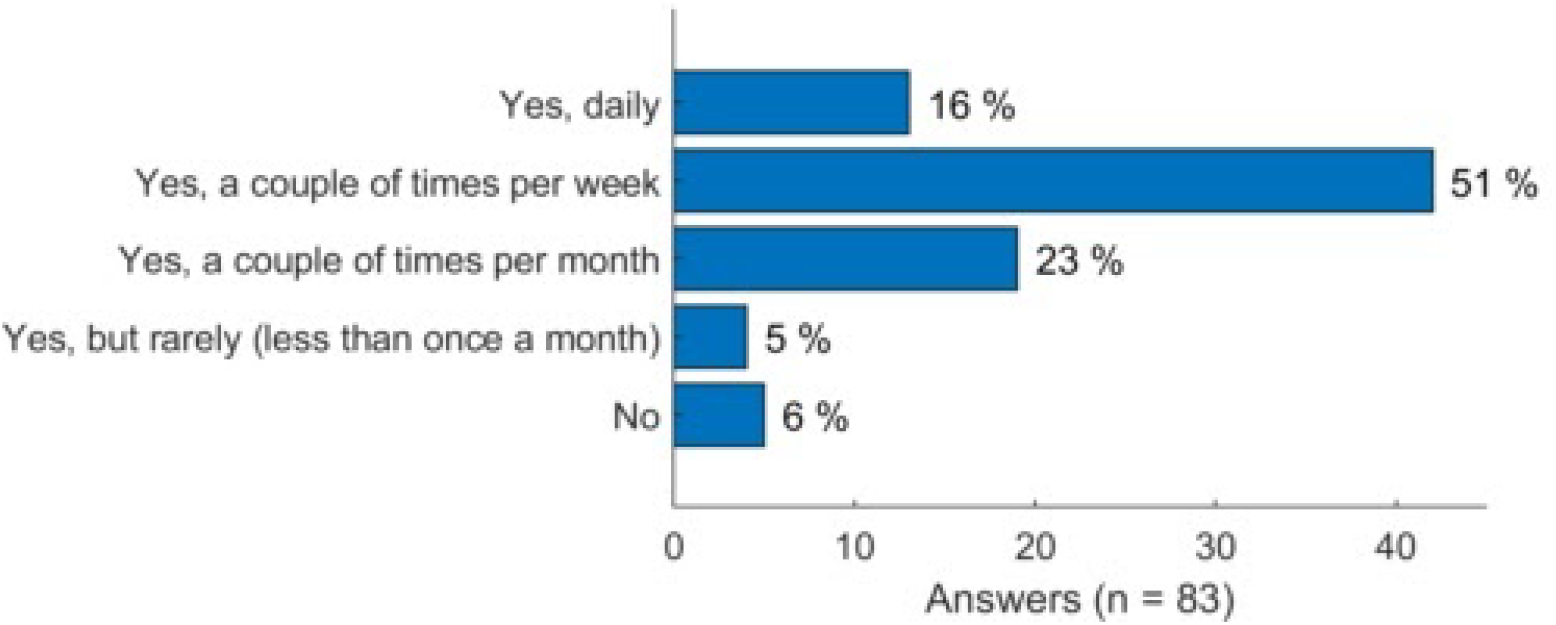

## Q7. Given the limited space for neuroscience in standards-based curricula, if you wanted to add neuroscience activities to your classroom, could you do it?

- Yes, I feel that neuroscience activities do meet our state standards
- Yes, I have independence on classroom activities, I can justify adding neuroscience even if not in our state standards
- Yes, I have time after our state standards exam to introduce new activities
- No, I cannot add activities unless they directly relate to our state standards
- Other

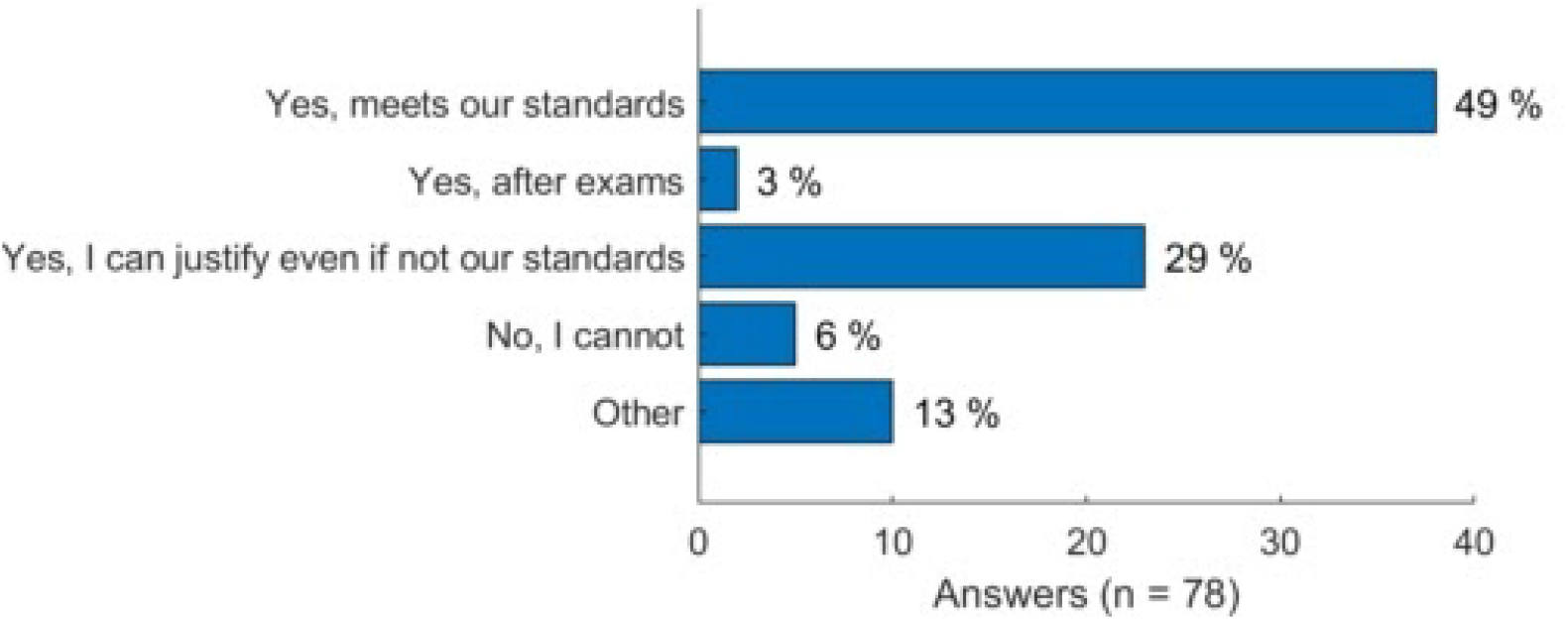

## Q8. Which of the following technologies are available to students taking your courses? (select all that apply)

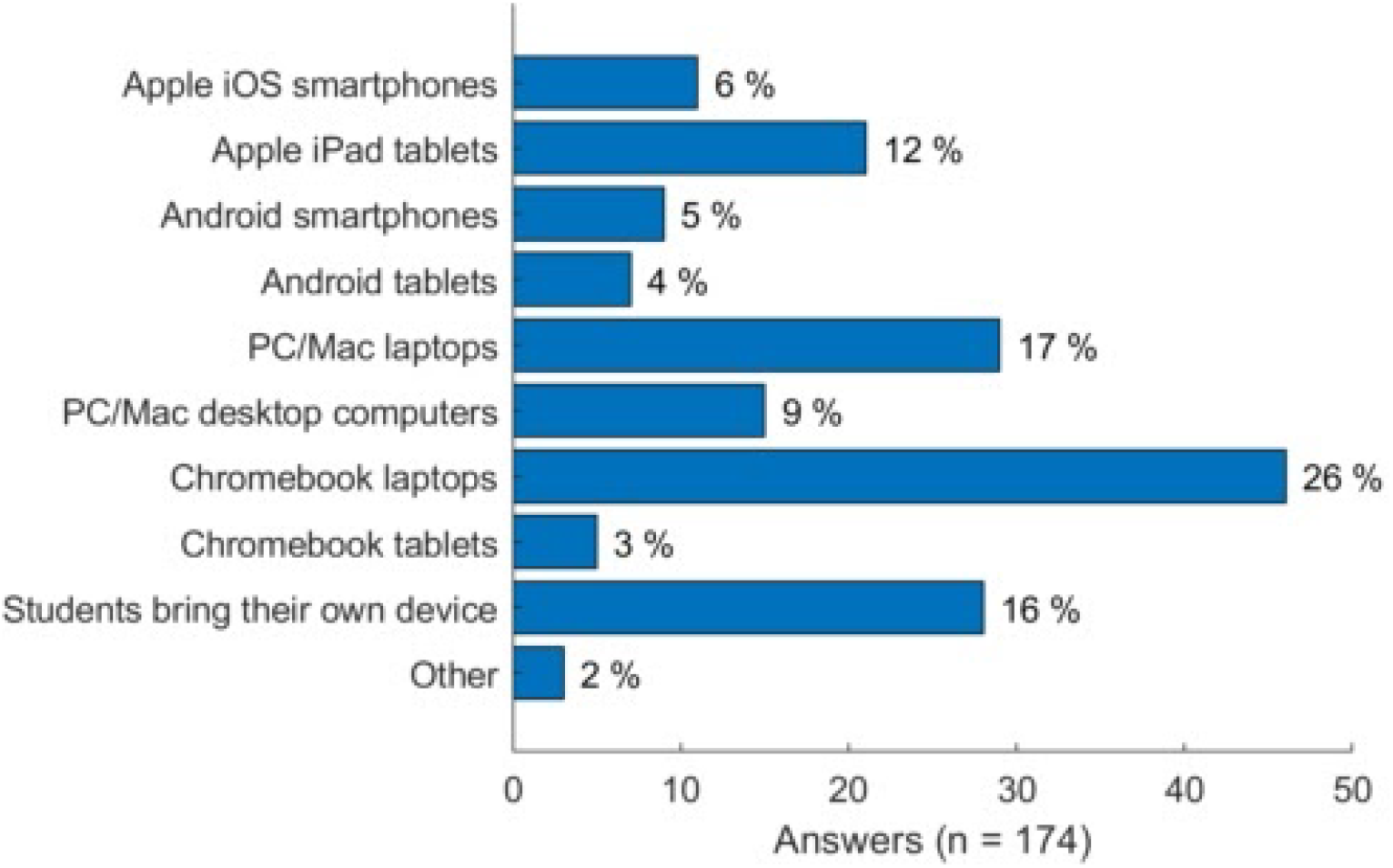

## Q9. Would you be interested in using SpikerBots to teach?

- Yes
- Maybe yes, but I don’t think this is sufficiently aligned with our standards
- No, I don’t think this is sufficiently aligned with our standards
- No, this looks too complicated / I don’t have time to learn this
- Other

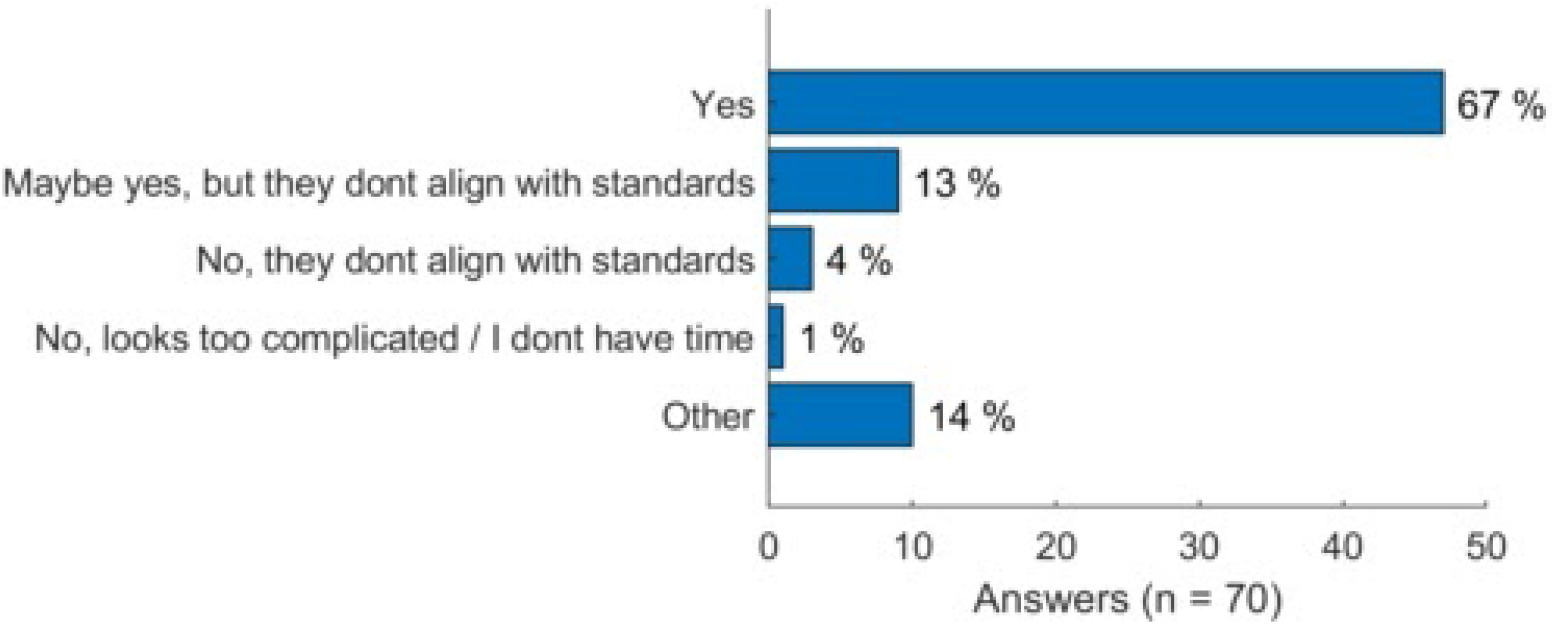

## Q10. Which types of instructional materials do you feel would most benefit students who are new to the SpikerBot neuroscience curriculum? (select up to 3 answers)

- Textbook
- Student worksheets (online & printable)
- Video library of demonstrations of laboratory exercises (ex-YouTube channel)
- STEM career connections (profession profiles, interviews with experts, networking opportunities, etc)
- Problem-based challenges and games (e.g. FIRST Robotics, FTC)
- Online platform to share brain designs

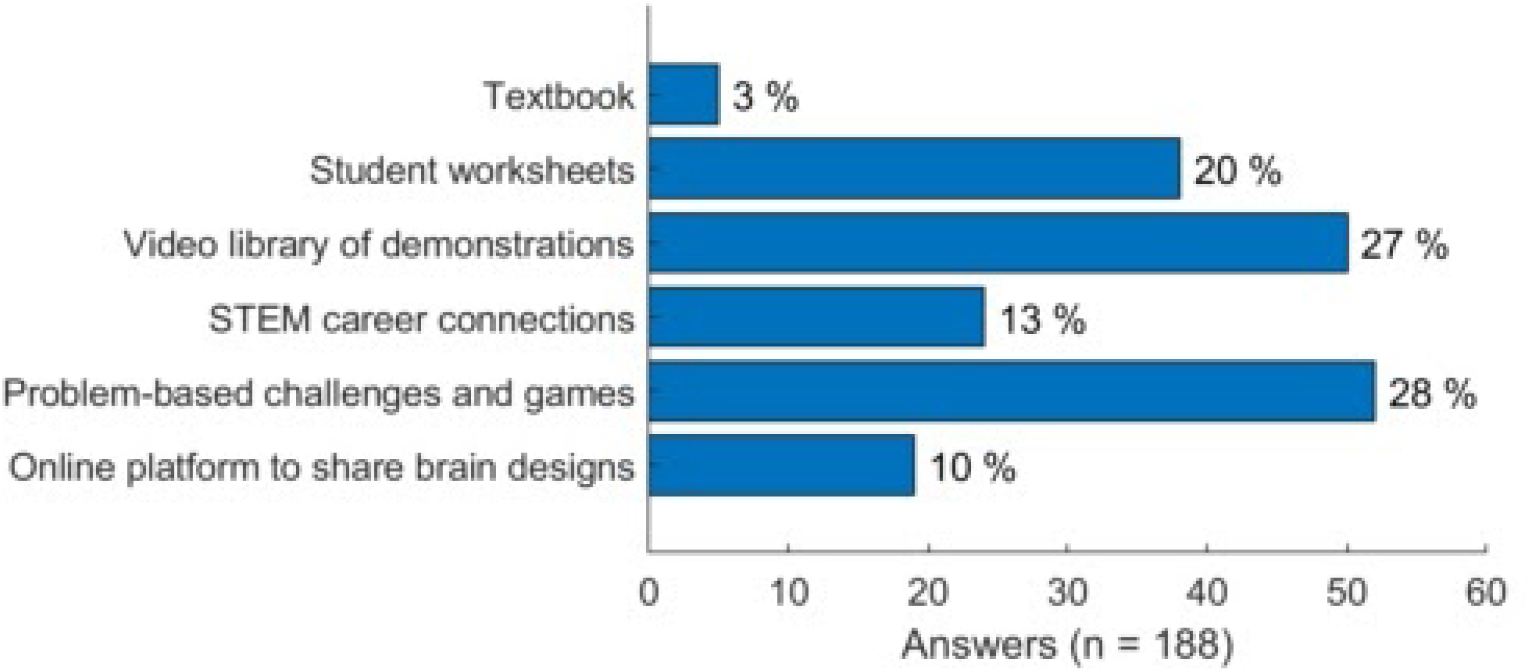

## Q11. How long do you think the SpikerBot-based curriculum should be (assuming each lesson is 60 minutes)?

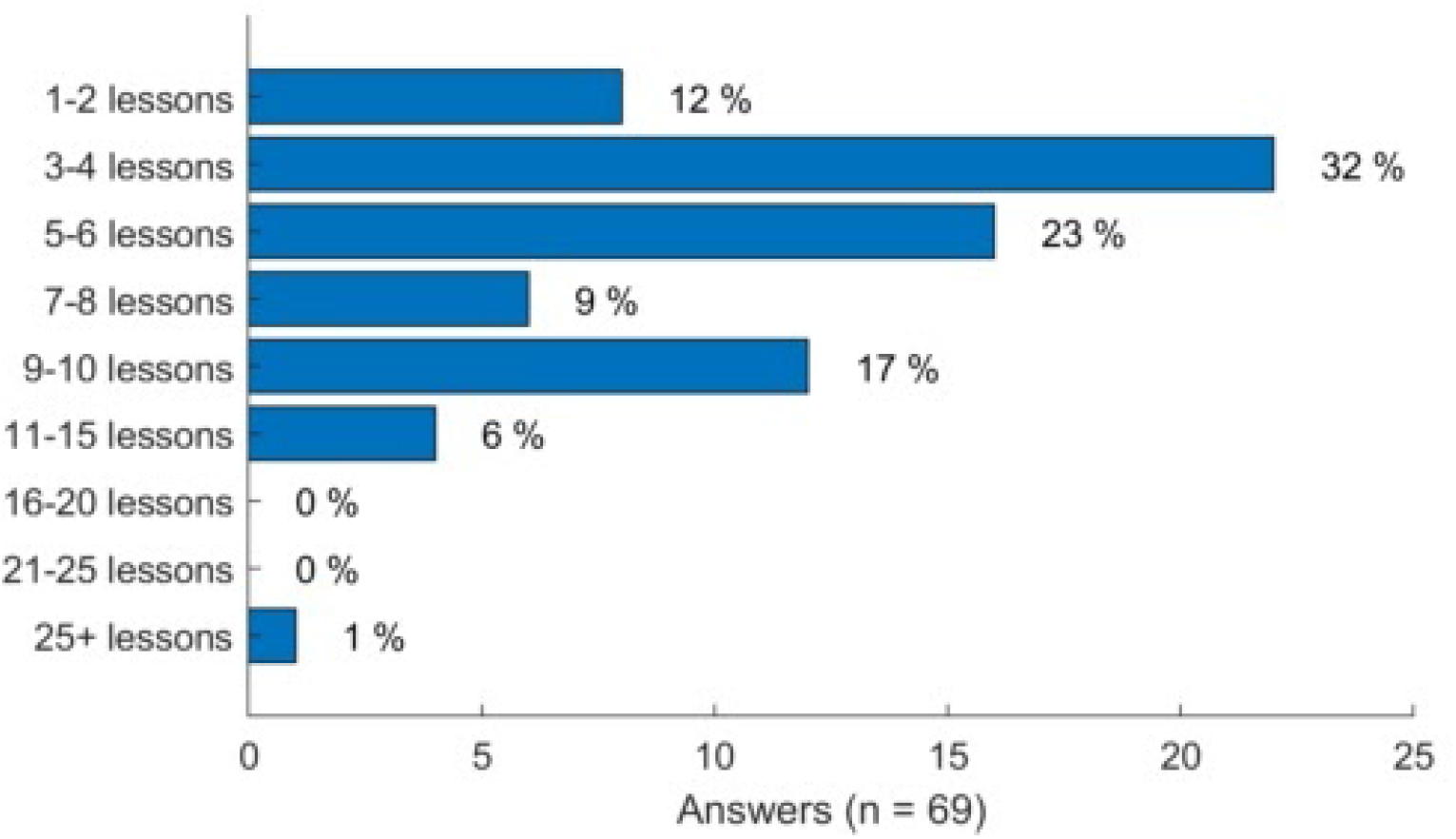

## Q12. If a classroom set of SpikerBots costs $2500 (about 10 robots for 30 students), how would you go about acquiring these funds? (select all that apply)

- I can use my department/school funds
- I would need additional private donations (ex. DonorsChoose)
- I would need to apply for a grant
- I would need to recruit other teachers
- Other, please specify

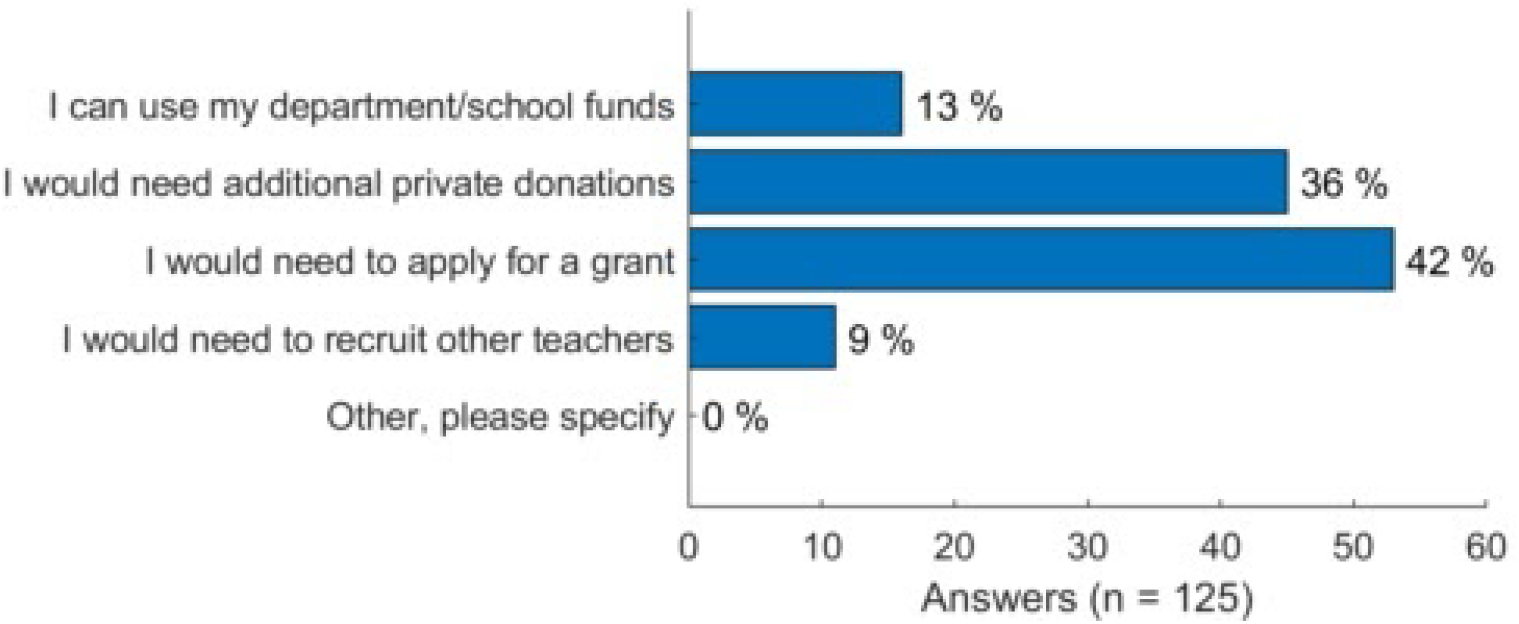

## Q13. What kind of professional development resources do you think would be most beneficial to teachers adopting SpikerBots for the first time? (select up to 3 answers)

- In person, half-day training workshop (post-COVID)
- In person, whole day training workshop (post-COVID)
- Live virtual training workshops
- Pre-recorded videos (ex-YouTube channel)
- In-class demonstrations by Backyard Brains
- Online teacher forum
- Online support by Backyard Brains
- Other

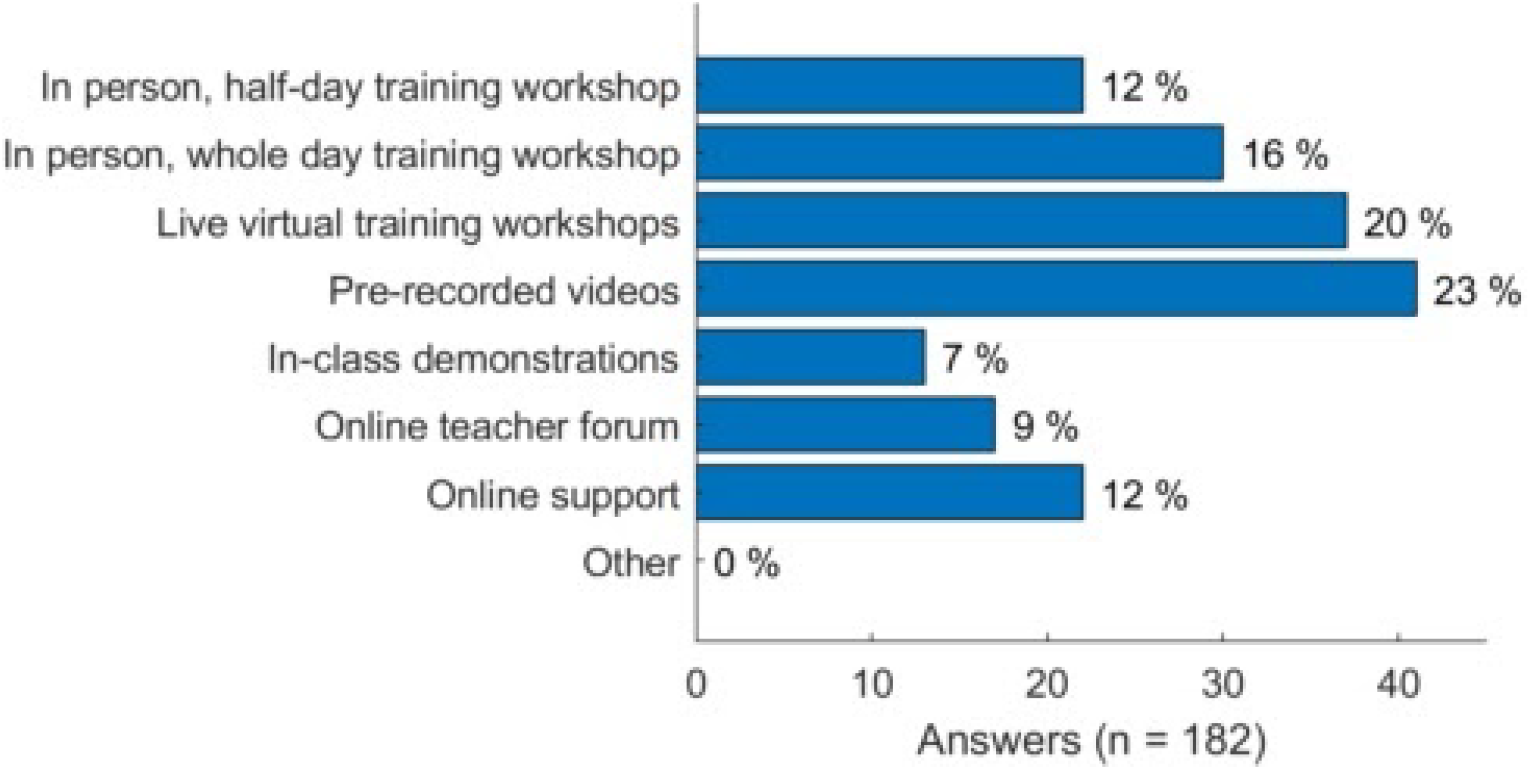

## Q14. Would learning to use the SpikerBot (ex. by participating in a workshop) count towards professional development hours/credits that are required by your school to collect?

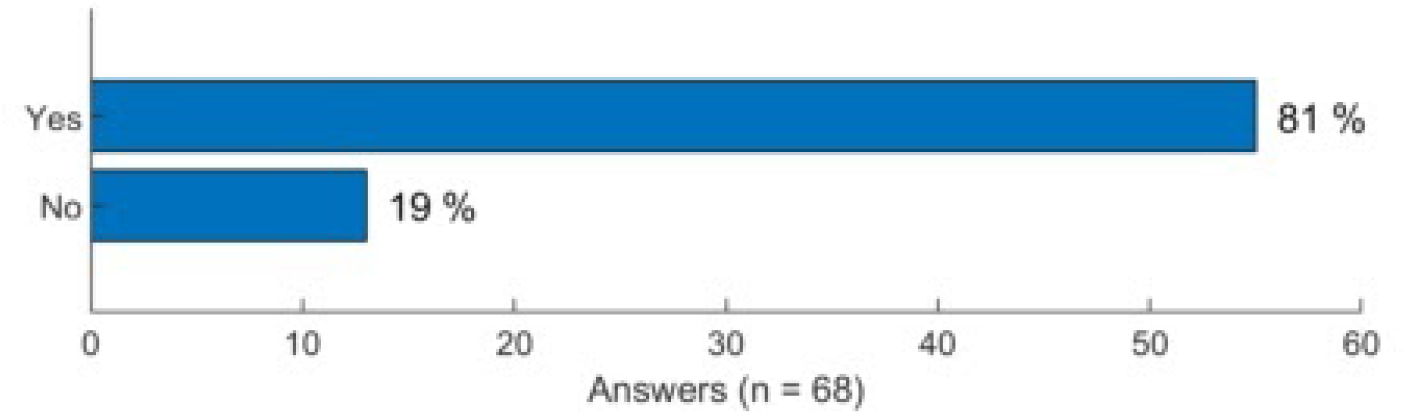

## Q15. How do you currently describe your gender?

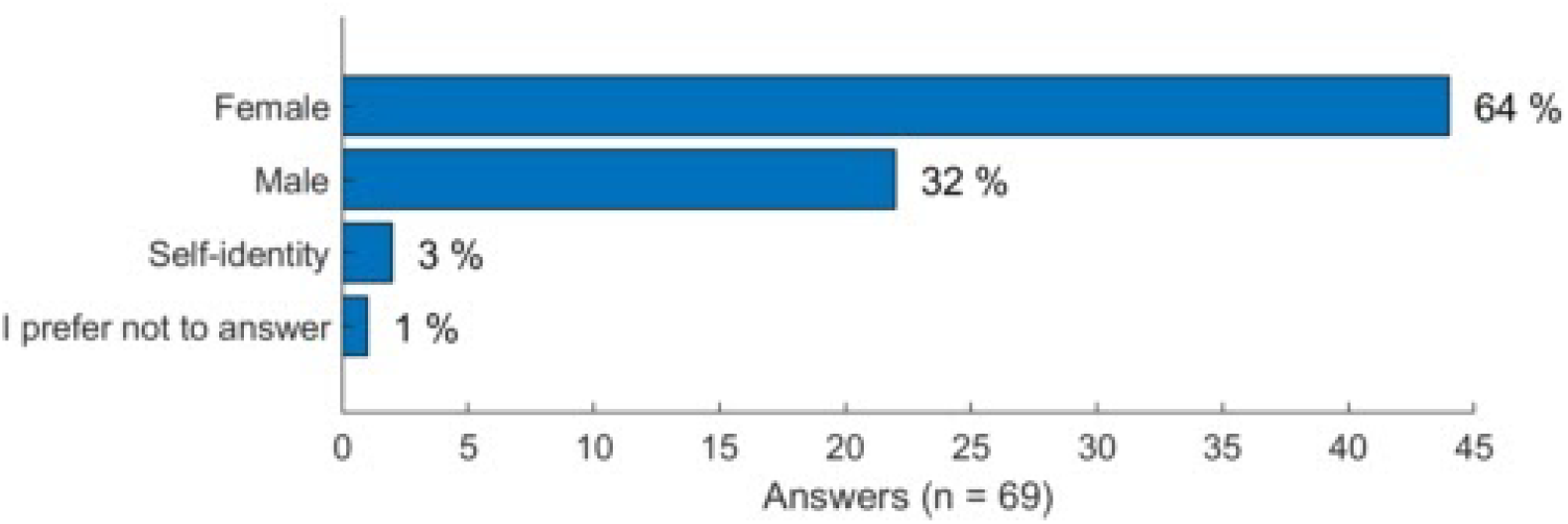

## Q16. How do you currently describe your race/ethnicity? (mark all options that correspond)

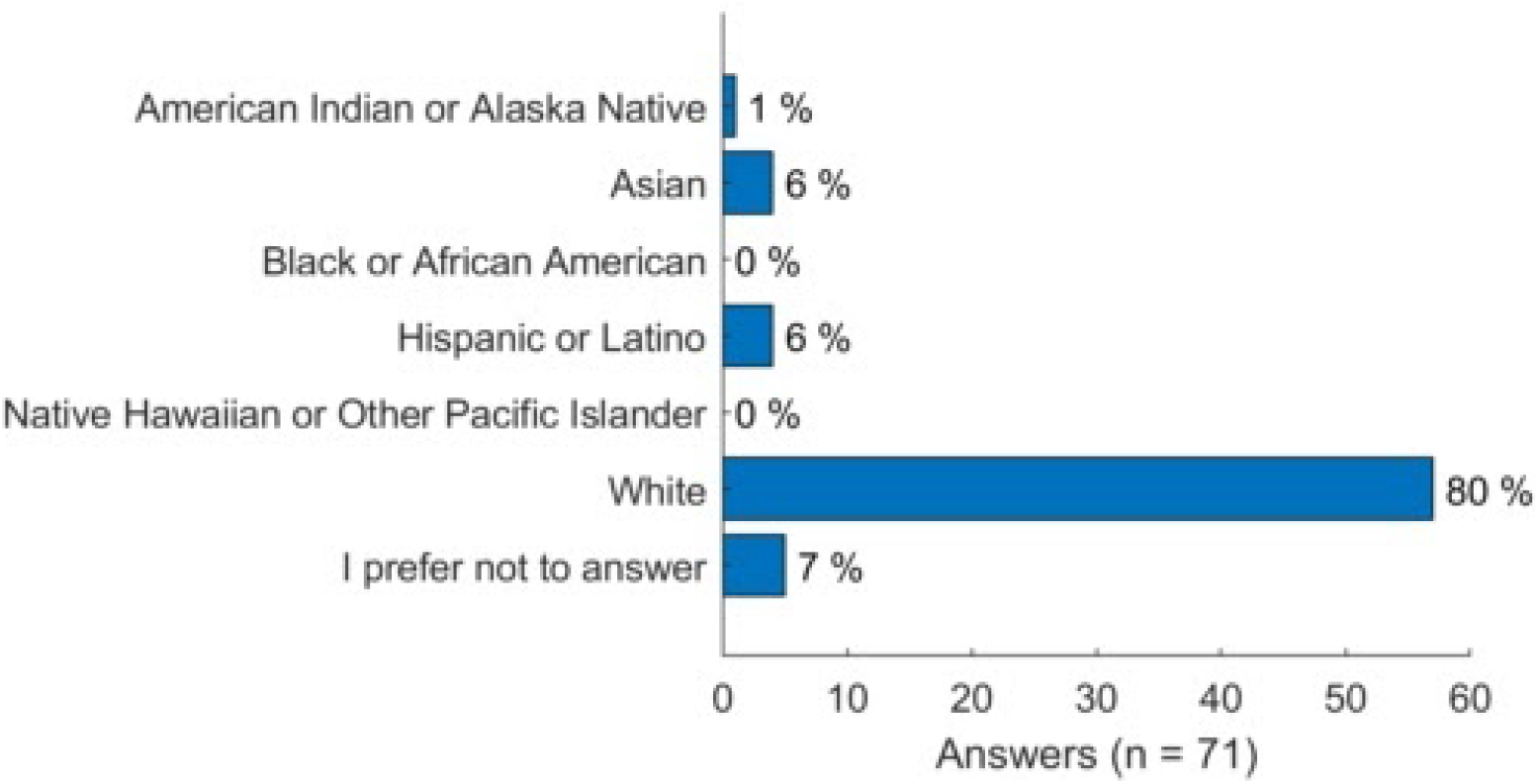

## Q17. How long have you been teaching?

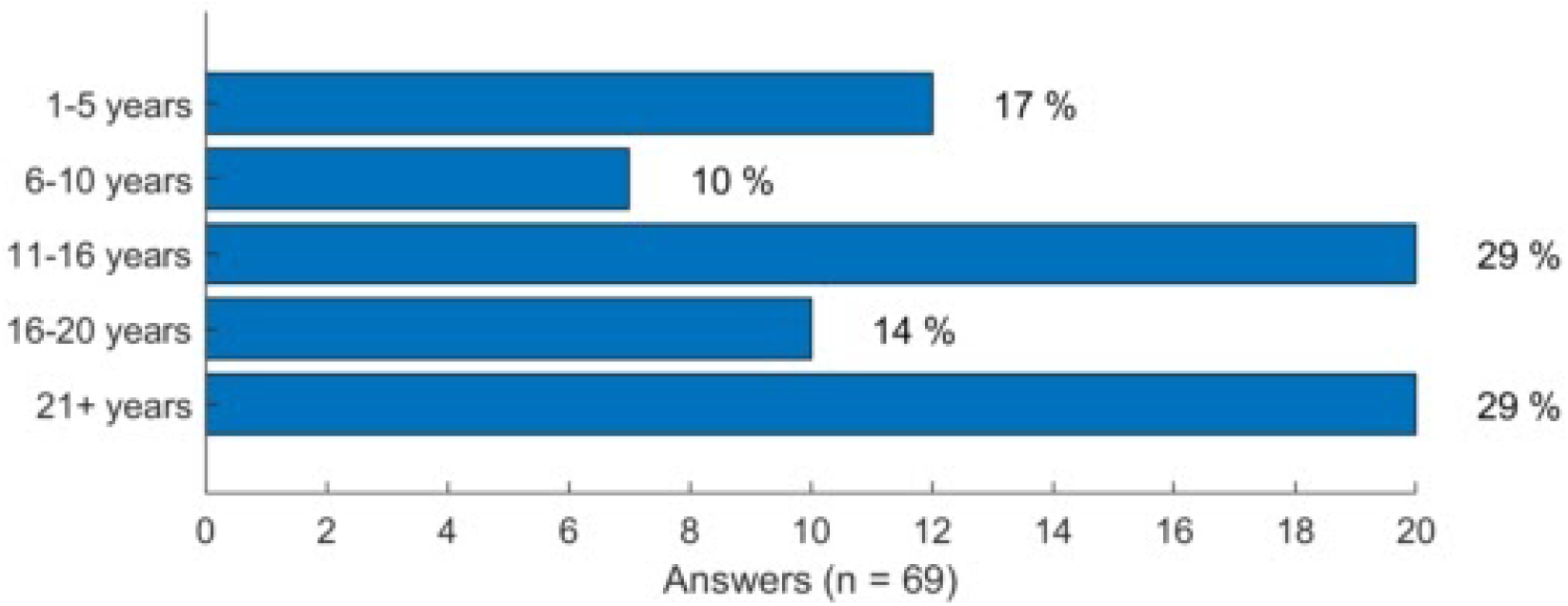

